# MHC class II antigen presentation by intestinal epithelial cells fine-tunes bacteria-reactive CD4 T cell responses

**DOI:** 10.1101/2023.01.23.525150

**Authors:** C. E. Heuberger, A. Janney, N. Ilott, A. Bertocchi, S. Pott, Y. Gu, M. Pohin, M. Friedrich, E. H. Mann, C. Pearson, F. M. Powrie, J. Pott, E. Thornton, K. J. Maloy

## Abstract

Although intestinal epithelial cells (IECs) can express major histocompatibility complex class II (MHC II), especially during intestinal inflammation, it remains unclear if antigen presentation by IECs favours pro- or anti-inflammatory CD4^+^ T cell responses. Using selective gene ablation of MHC II in IECs and IEC organoid cultures, we assessed the impact of MHC II expression by IECs on CD4^+^ T cell responses and disease outcomes in response to enteric bacterial pathogens. We found that intestinal bacterial infections elicit inflammatory cues that greatly increase expression of MHC II processing and presentation molecules in colonic IECs. Whilst IEC MHC II expression had little impact on disease severity following *Citrobacter rodentium* or *Helicobacter hepaticus* infection, using a colonic IEC organoid-CD4^+^ T cell co-culture system, we demonstrate that IECs can activate antigen-specific CD4^+^ T cells in an MHC II-dependent manner, modulating both regulatory and effector Th cell subsets. Furthermore, we assessed adoptively transferred *H. hepaticus*-specific CD4^+^ T cells during intestinal inflammation *in vivo* and report that IEC MHC II expression dampens pro-inflammatory effector Th cells. Our findings indicate that IECs can function as non-conventional antigen presenting cells and that IEC MHC II expression fine-tunes local effector CD4^+^ T cell responses during intestinal inflammation.

## Introduction

There is increasing appreciation of the important roles that tissue-resident cells play in local immune responses, particularly at barrier surfaces, such as the gut, skin, and lung^1,2^. In the gut, a single cell layer of intestinal epithelial cells (IECs) forms a physical barrier between the luminal microbiota and the leucocytes within the underlying lamina propria. Sensing of microbes by IECs induces innate defence mechanisms and helps maintain barrier function^3,4^. In addition, IECs have long been known to express major histocompatibility complex class II (MHC II) molecules and associated processing molecules, especially during intestinal inflammation, suggesting that they may influence antigen-specific immune responses by intestinal CD4^+^ T cells^5–9^. MHC II expression by IECs is compartmentalized along the gut, with small intestinal IECs constitutively expressing high levels^7,9–12^, whereas colonic IECs do not express MHC II molecules under steady state conditions^13^. However, in rodent models of intestinal inflammation, MHC II expression is upregulated in both small intestinal and colonic IECs^14–16^, and IECs from inflammatory bowel disease (IBD) patients also express MHC II during active disease^13,17–19^.

The functional impact of IEC MHC II expression on local CD4^+^ T cell responses remains uncertain, with different effects reported by distinct studies. Several studies have provided evidence that IEC MHC II expression may potentiate pathogenic CD4^+^ T cell responses in the inflamed gut. For example, MHC II expressing IECs isolated from IBD patients induced the proliferation of CD4^+^ T cells and the release of IFN-γ *ex vivo*^18^. Similarly, it was reported that mice selectively lacking MHC II expression in IECs exhibited reduced severity in models of colitis induced by DSS administration or CD4^+^ T cell transfer^20^. Furthermore, it was recently reported that small intestinal IEC MHC II expression was both necessary and sufficient for activation of pathogenic donor CD4^+^ T cells that mediate enteritis during experimental graft-versus-host disease (GvHD)^16^.

Conversely, other studies have proposed that IEC MHC II expression acts to boost immune-regulatory responses. For example, autoantigen expression by IECs was reported to drive the expansion of antigen-specific CD4^+^ FoxP3^+^ regulatory T cells (Tregs)^21^. Another study found that transgenic mice lacking MHC II expression in all non-haematopoietic cells (including IECs) developed more severe CD4^+^ T-cell-mediated colitis than wild-type littermates^22^. In addition, a recent study reported that small intestinal IEC MHC II expression supported the diurnal expression of IL-10 by intraepithelial lymphocytes^23^. These conflicting results regarding the ability of IECs to preferentially promote effector or regulatory T cell responses suggest context- and location-specific effects and highlight the need for further investigation. In particular, a better understanding of the effects of IEC MHC II expression on local CD4^+^ T cell responses during intestinal inflammation could reveal new aspects of IBD pathogenesis.

We therefore generated transgenic mice that selectively lacked MHC II on IECs (IEC^ΔMHCII^) and subjected them to well characterized models of bacterially-driven intestinal inflammation. In parallel, we utilized primary IEC organoids^24^ to develop an *in vitro* co-culture assay to investigate the impact of IEC MHC II expression specifically on bacteria-specific CD4^+^ T cells in a well-controlled setting. We report that IECs respond to inflammatory mediators with increased expression of MHC II processing and presentation genes, that IECs take up and process antigen and activate CD4^+^ T cells in an MHC II-dependent manner. *In vivo*, we show that IEC-specific deletion of MHC II does not impair induction of effector CD4^+^ T cell responses and has no significant impact on intestinal inflammation or host survival following infection with the attaching-effacing (A/E) pathogen *Citrobacter rodentium*. Similarly, we report that ablation of MHC II expression by IECs has little impact on CD4^+^ T cell-mediated chronic typhlocolitis triggered by infection with *Helicobacter hepaticus* and IL-10R blockade^25^. By contrast, during *H. hepaticus*-driven colitis, we observed that IEC MHC II dampened bacteria-specific CD4^+^ T cell pro-inflammatory cytokine expression. Together, our results demonstrate that while not essential for priming of CD4^+^ T cell responses during bacterial infections, MHC II antigen-presentation by IECs fine-tunes the functional capabilities of effector CD4^+^ T cells in the intestine to potentially favour restoration of tissue homeostasis.

## Results

### Primary colonic IECs increase expression of MHC II processing and presentation molecules in response to inflammatory cues

To analyse the response of IECs to intestinal inflammation, we grew colonic IEC organoids from intestinal stem cells of steady state mice. To capture the intestinal inflammatory milieu, we isolated lamina propria leucocytes (LPLs) from colitic mice (infected with *Helicobacter hepaticus* + anti-IL10R mAb treatment weekly for 2 weeks^25^) and cultured them overnight to produce inflammatory conditioned medium (iCM). We obtained control conditioned medium (cCM) from LPLs isolated and cultured from untreated mice. LPL cell numbers were normalised to produce cCM and iCM. We then stimulated primary colonic IEC organoids with 10% iCM or cCM (**Figure 1A**). To obtain a global transcriptional response of primary IECs, we performed bulk RNA sequencing (RNA-Seq) from iCM- and cCM-stimulated organoids. A large number of genes were differentially expressed in IECs in response to iCM stimulation (**Table S1**), with H2-Ab1 among the top 10 differentially expressed genes (**Figure 1B**). We also found that iCM-treated primary IEC organoids exhibited significantly increased expression of IFN-γ response genes (*Itgp*), chemokines *(Cxcl11)*, as well as co-inhibitory (*Cd274*) molecules (**Figure 1C**). A gene set enrichment analysis on Gene Ontology categories indicated an increased expression of genes related to antigen processing and cross presentation in IECs treated with iCM (**Figure 1D**). Genes associated with the antigen presentation pathway included cathepsins (*Ctss* and *Ctsb*) as well as the MHC II associated invariant chain (*Cd74*), among others (**Figure 1E**). As IFN-γ signalling was among the top 10 significantly upregulated pathways, we assessed the functional response of IECs to iCM stimulation and detected the rapid phosphorylation of STAT1, while STAT3 was phosphorylated in response to both iCM and cCM (**Figure S1A**). In addition, the expression of IFN-γ response genes (*Igtp* and *Isg15*) were induced in organoids after 20h of stimulation with iCM, while *Reg3g* (IL-17 response gene) and *Cxcl2* (general inflammatory response gene) expression was induced in response to both iCM and cCM stimulation (**Figure S1B-E**). These results indicate that primary colonic IECs respond strongly to inflammatory mediators and that this includes upregulation of antigen processing and presentation capabilities.

**Figure 1:**
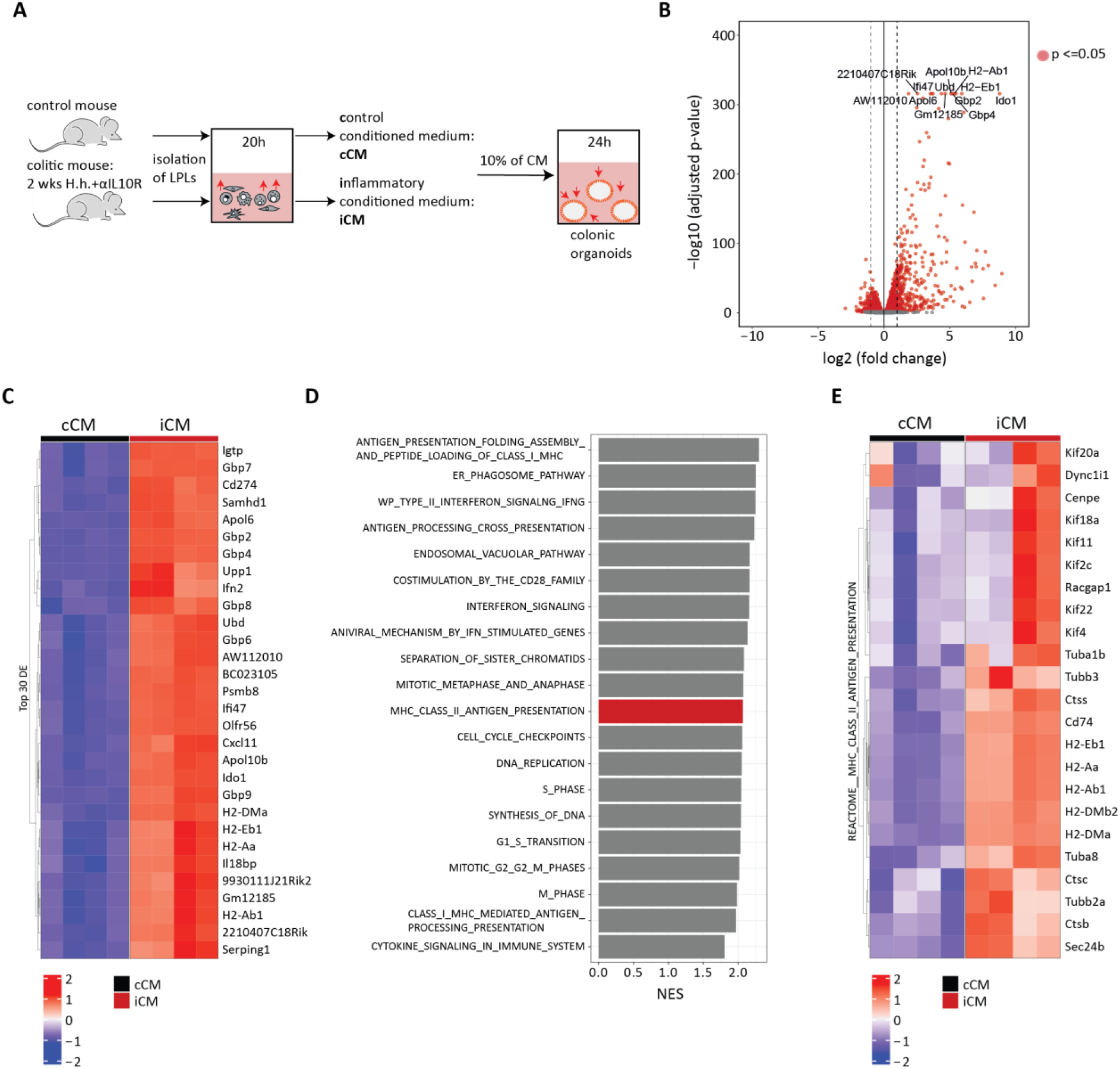
Primary colonic IECs upregulate expression of MHC II processing and presentation molecules in response to inflammatory cues. (A-E) RNA-seq was performed on colonic organoids stimulated with 10% iCM or cCM for 24h. (A) Schematic of experimental setup. (B, C) Differentially expressed genes were calculated using DESeq2 to determine the FDR. Genes were considered differentially expressed at an adjusted p-value <0.05. (B) Volcano plot of significantly differentially expressed genes. The top ten of differentially expressed genes are highlighted. (C) List of the top 30 of significantly differentially expressed genes. (D) Significantly enriched disease pathways in the set of genes significantly upregulated in iCM compared to cCM treated organoids. Gene sets were considered significantly enriched at an adjusted p-value <0.05. (E) Heatmap depicting antigen processing and presentation associated genes differentially expressed and significantly upregulated in iCM vs cCM treated organoids. H.h. – *Helicobacter hepaticus*, iCM – inflammatory conditioned medium, cCM – control conditioned medium, LPLs – lamina propria lymphocytes, NES – normalised enrichment score.

### IFN-γ plays a key role in inducing MHC II expression in primary colonic IECs

To analyse the potential functional impact of MHC II by IECs on intestinal inflammation *in vivo*, we generated transgenic mice lacking H2-Ab1 in IECs, by crossing H2-Ab1^fl/fl^ mice^26^ with mice expressing Cre recombinase under the control of the Villin promoter^27^ resulting in H2-Ab1^VC^ (hereafter called IEC^ΔMHCII^) or H2-Ab1^fl/fl^ (hereafter called IEC^WT^) mice. By using breeders that were heterozygous for the *Cre* allele, we generated littermate controls (IEC^WT^) that were co-housed with their IEC^ΔMHCII^ littermates during all experiments, to prevent any potential differences in the microbiota. In addition, as there have been reports that the Villin-Cre strain can be ‘leaky’, resulting in off-target deletion of the floxed alleles^28^, we validated each individual IEC^ΔMHCII^ mouse by flow cytometry to verify selective ablation of MHC II in IECs, with maintenance of MHC II expression in haematopoietic lineages **(Figure S2A and B)**.

We first wanted to identify the inflammatory cytokine(s) responsible for the induction of MHC II expression by primary colonic IECs. We used a panel of cytokines that have been implicated in intestinal inflammation (IFN-γ, IL-22, IL-17a, TNFα, IL-13, IL-18, IFN-λ2, IFN-β, IL-6 and IL-25) to stimulate primary colonic IEC organoids and found that only IFN-γ could induce MHC II mRNA expression (**Figure 2A**). As IFN-γ induces MHC II expression in many cell types through activation of the class II major histocompatibility complex transactivator (CIITA)^29^, we investigated the kinetics of MHC II and CIITA mRNA expression in IFN-γ stimulated primary colonic IEC organoids. While MHC II mRNA expression increased after 6h and peaked at 8h (**Figure 2B**), CIITA mRNA expression was already upregulated after 2h (**Figure 2C**).

**Figure 2:**
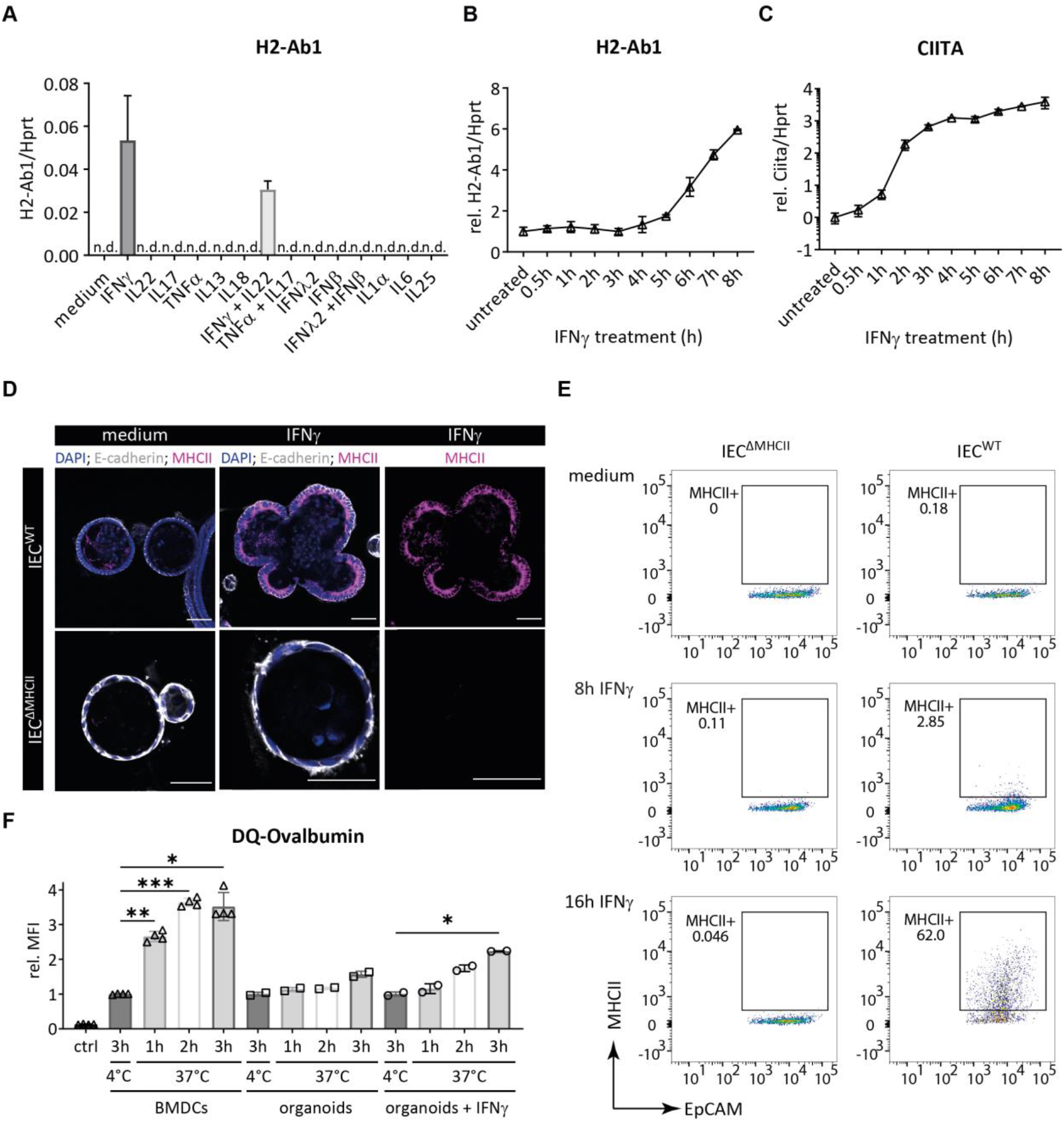
IECs express MHC II in response to IFN-γ and take up and process antigen. (A) IEC^WT^ organoids were stimulated with the indicated cytokines (20ng/ml) for 16h and *H2-Ab1* expression was assessed by qPCR analysis. (B) IEC^WT^ organoids were stimulated with IFN-γ (20ng/ml) for the indicated amount of time and *H2-Ab1* expression was assessed by qPCR analysis. (C) IEC^WT^ organoids were stimulated with IFN-γ (20ng/ml) for the indicated amount of time and *Ciita* expression was assessed by qPCR analysis. (D) IEC^WT^ and IEC^ΔMHCII^ organoids were stimulated with IFN-γ (20ng/ml) for 16h and MHC II expression was assessed by immunofluorescent imaging. Organoids were stained for MHC II (magenta), E-cadherin (grey) and nuclei with DAPI (blue). (E) IEC^WT^ and IEC^ΔMHCII^ organoids were stimulated with IFN-γ (20ng/ml) or left untreated for the indicated amount of time and MHC II expression was assessed by flow cytometry. (F) BMDCs and IEC^WT^ organoids were stimulated with DQ-Ovalbumin for the indicated times and at the indicated temperatures. IEC^WT^ organoids were pre-treated with IFN-γ (20ng/ml) for 16h prior to DQ-Ovalbumin incubation where indicated. The relative median fluorescent intensity (MFI) was assessed by flow cytometry. Data are representative from one of two independent experiments, with two technical replicates run in duplicates (A-C and F). Data are representative from one of two independent experiments, with two technical replicates (D and E). Bars and symbols denote mean (± SD) (A-C) or mean (±SEM) as a relative to 3h incubation at 4°C (F). Scale bars, 50μm. Statistical significance between the groups was determined by mixed-effects analysis with Geisser-Greenhouse correction and Tukey’s multiple comparisons test (F).

We performed confocal microscopy to validate MHC II protein expression. Colonic IEC^WT^ organoids exhibited prominent cytosolic and surface protein expression of MHC II following stimulation with IFN-γ, whereas this was undetectable on colonic IEC^ΔMHCII^ organoids (**Figure 2D**). These findings were confirmed by flow cytometry, where we found that a high proportion of wild type (IEC^WT^) colonic IEC expressed MHC II after 16h of IFN-γ stimulation (**Figure 2E**). In contrast, colonic IECs from IEC^ΔMHCII^ mice did not express MHC II protein in response to IFN-γ stimulation (**Figure 2E**). Another important pre-requisite to function as antigen presenting cells (APCs) is the ability to take up and process antigen. We used DQ-Ovalbumin^30^, a self-quenched conjugate of ovalbumin that fluoresces upon proteolytic degradation, to assess the capability of IECs to take up and process antigen. We detected a significant increase in DQ-Ovalbumin MFI after 3h of DQ-Ovalbumin incubation when primary colonic IEC organoids had been pre-stimulated with IFN-γ (**Figure 2F**). Together, these findings highlight that, during inflammatory conditions, colonic IECs have the potential to function as non-conventional APCs.

### Colonic IEC MHC II expression does not alter disease severity driven by bacterial infection

We next assessed the role of MHC II expression by colonic IECs in intestinal infection and inflammation models. Following oral infection of IEC^ΔMHCII^ and IEC^WT^ littermates with *C. rodentium*, mice were sacrificed at two weeks post-infection (**Figure 3A)**. Flow cytometric analyses revealed that *C. rodentium* infection induced MHC II expression in approximately 20% of colonic IECs in IEC^WT^ mice, whereas MHC II remained undetectable on IECs from IEC^ΔMHCII^ mice (**Figure 3B**). Assessment of the body weight throughout the course of *C. rodentium* infection did not reveal significant differences between IEC^WT^ and IEC^ΔMHCII^ littermates (**Figure 3C**). Moreover, we found that *C. rodentium* infection induced comparable levels of intestinal inflammation, as determined by histological scoring in the colon and caecum of IEC^WT^ and IEC^ΔMHCII^ littermates (**Figure 3D-F**), which was in accordance with similar bacterial colonisation levels (**Figure 3G-I**). Our findings indicate that MHC II expression on IECs is dispensable for both intestinal inflammation and for control and survival of *C. rodentium* infection.

**Figure 3:**
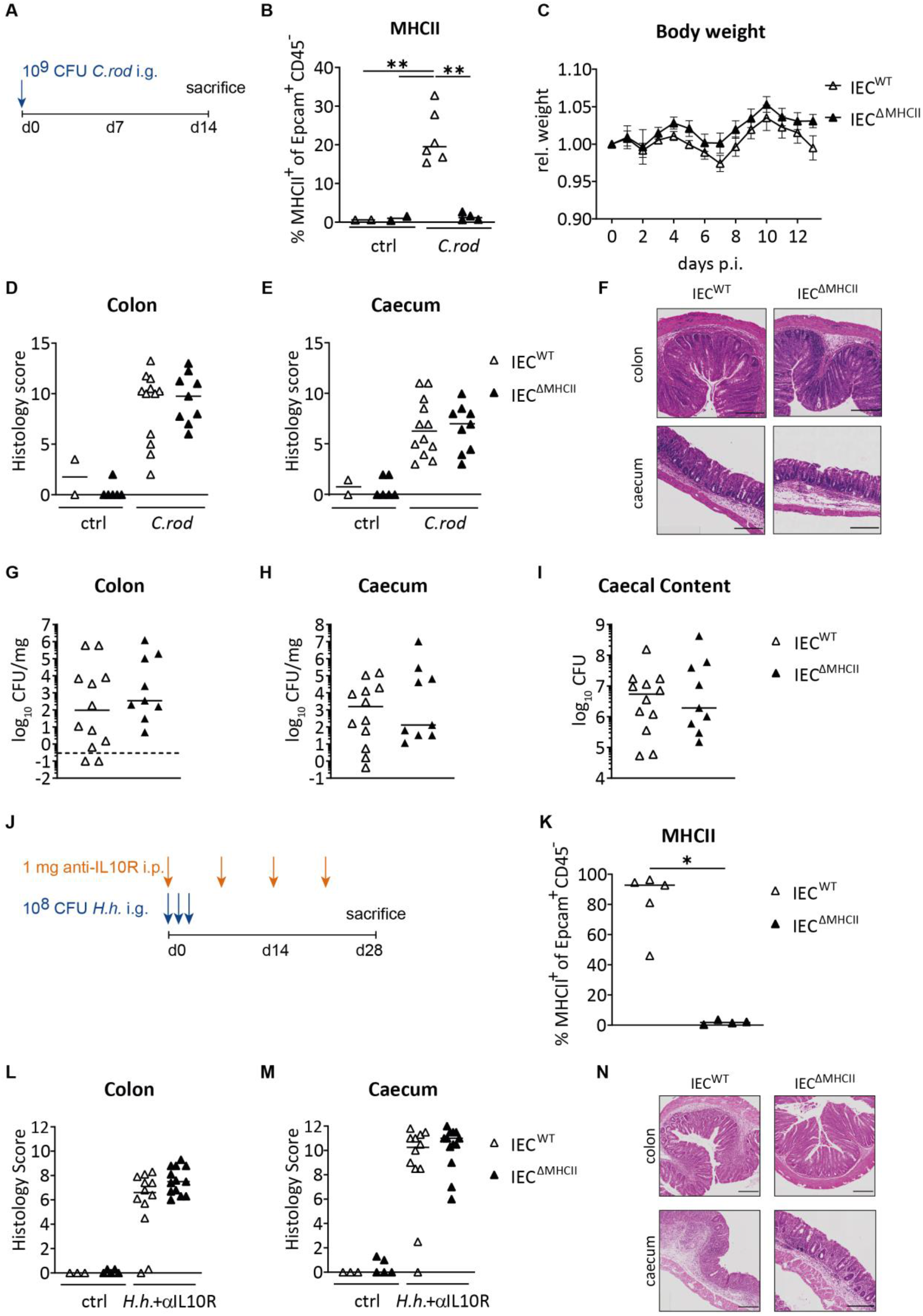
IEC MHC II expression plays a negligible role in bacterial driven colitis. (A-I) IEC^WT^ and IEC^ΔMHCII^ littermates were orally infected with 10^9^ CFU (colony-forming units) *C. rodentium*. (A) Schematic of treatment protocol. (B) Colonic IEC MHC II expression of IEC^WT^ and IEC^ΔMHCII^ littermates was assessed by flow cytometry. (C) Weight curves of IEC^WT^ and IEC^ΔMHCII^ littermates during acute colitis. (D and E) Histopathology of the colon (D) and caecum (E) was assessed on day 14. (F) Representative hematoxylin and eosin (H&E) sections of colon (top row) and caecum (bottom row) of treated IEC^WT^ and IEC^ΔMHCII^ littermates at day 14. (G-I) Tissue adherent CFU in colon (G), caecal tissue (H) and planktonic CFU in caecal content (I) of IEC^WT^ and IEC^ΔMHCII^ littermates at day 14 p.i.. (J-N) IEC^WT^ and IEC^ΔMHCII^ littermates were orally infected with 10^8^ CFU *H. hepaticus* on three consecutive days and injected with 1mg anti-IL10R weekly. (J) Schematic of treatment protocol. (K) Colonic IEC MHC II expression of IEC^WT^ and IEC^ΔMHCII^ littermates was assessed by flow cytometry. (L and M) Histopathology of the colon (L) and caecum (M) was assessed on day 28. (N) Representative H&E sections of colon (top row) and caecum (bottom row) of treated IEC^WT^ and IEC^ΔMHCII^ littermates at day 28. Data from one experiment of n=4 IEC^ΔMHCII^ and n=6 IEC^WT^ infected mice and n=2 IEC^ΔMHCII^ and n=2 IEC^WT^ uninfected mice (B). Data from two pooled independent experiments with total mouse numbers per group as follows; n=9 IEC^ΔMHCII^ and n=12 IEC^WT^ infected mice and n=6 IEC^ΔMHCII^ and n=2 IEC^WT^ uninfected mice (C-I). Data from one experiment with total mouse numbers per group as follows; n=4 IEC^ΔMHCII^ and n=5 IEC^WT^ treated mice (K). Data from two pooled independent experiments with total mouse numbers per group as follows; n=13 IEC^ΔMHCII^ and n=12 IEC^WT^ treated mice and n=5 IEC^ΔMHCII^ and n=3 IEC^WT^ untreated mice (L-N). Symbols denote individual mice (B and D-M) or mean (± SEM) (C). Horizontal bars indicate group medians. Dashed horizontal line depicts theoretical detection limit of assay. Scale bars, 200*μ*m. Statistical significance between the groups was determined by two-way ANOVA multiple comparisons test with Tukey’s correction (B) or Sidak’s correction (C), or Mann-Whitney test (D-M). P-value summary: *p<0.05; **p<0.01. CFU – colony-forming units, ctrl – control, *C.rod* – *Citrobacter rodentium*, *H.h*. – *Helicobacter hepaticus*, p.i. – post infection.

We also assessed the importance of IEC MHC II expression in a model of chronic intestinal inflammation induced by oral infection with *H. hepaticus* and blockade of the regulatory response with anti-IL10R antibody (Hh+αIL10R)^25^ (**Figure 3J**). Flow cytometric analyses revealed that Hh+αIL10R treatment induced MHC II expression in approximately 90% of colonic IECs in IEC^WT^ controls, whereas MHC II remained undetectable on IECs from IEC^ΔMHCII^ mice (**Figure 3K and S2C**).

Four weeks after disease induction, histological assessment of inflammation in the colon and caecum revealed comparable disease severity between IEC^ΔMHCII^ and IEC^WT^ littermates (**Figure 3L-N**). As inflammation in this model is driven by pathogenic Th1 and Th17 cell responses^25^, we assessed frequencies of CD4^+^ T cells and Treg subsets in the colonic LP of Hh+αIL10R treated IEC^ΔMHCII^ mice and IEC^WT^ littermates and found no significant differences (**Figure S2D-H)**. Taken together, these data reveal that MHC II expression by colonic IECs is dispensable for *H. hepaticus* triggered chronic colitis and has no impact on disease severity.

### IECs induce antigen-specific activation of CD4^+^ T cells via MHC II *ex vivo*

The potent inflammatory responses induced in the bacterial infection models may have masked more subtle effects of antigen presentation by IECs. Therefore, to assess whether IECs can function as non-conventional APCs to modulate CD4^+^ T cells, we established a co-culture system comprising colonic IEC organoids and TCR transgenic CD4^+^ T cells. We used *H. hepaticus-specific* CD4^+^ T cells (HH7-2tg-FoxP3^hCD2^ cells), which express a TCR specific for the *H. hepaticus-unique* peptide HH_1713^31^ and allow the identification of FoxP3^+^ Tregs using the surface marker hCD2. As IECs primarily encounter primed effector T cells in the gut, we pre-activated the HH7-2tg-FoxP3^hCD2^ cells *in vitro* (3 days anti-CD3/anti-CD28 and 2 days rest) before adding them to colonic IEC organoids in the presence or absence of the cognate peptide HH_1713. To induce MHC II expression, colonic IEC organoids were pre-treated with IFN-γ in some culture conditions. We assessed expression of canonical transcription factors (TF) that drive Th1/Th17 effector and regulatory T cell subsets in HH7-2tg cells after co-culture with colonic IEC organoids. We detected a significant increase in FoxP3^+^, RORγt^+^ and Tbet^+^ (**Figure 4A-F**) HH7-2tg-FoxP3^hCD2^ T cells when they were able to interact with the antigen presented on IFN-γ-stimulated IECs. TF upregulation by HH7-2tg-FoxP3^hCD2^ cells was dependent on MHC II expression by IECs, as there was no response when IEC^ΔMHCII^ colonic IEC organoids were used in the co-cultures (data not shown), or when MHC II was blocked with a blocking antibody (**Figure 4A-F**). Thus, MHC II antigen presentation by colonic IECs enhances both regulatory and effector Th cell subsets. We also assessed cytokine expression after 3 days of co-culture and detected significant induction of TNF-α expression by HH7-2tg-FoxP3^hCD2^ cells, which was again dependent on MHC II presentation of the cognate peptide by IFN-γ-stimulated IECs (**Figures 4G and H**). In addition, under these conditions, we detected significant production of IFN-γ, CXCL-10 and GM-CSF but not of TNF-α, IL-17A, Granzyme B, IL-10 or IL-4 in the co-culture supernatants (**Figure 4I**). These observations indicate that MHC II presentation of cognate peptide by primary IECs can modulate CD4^+^ effector T cell functions. Furthermore, we observed a marked upregulation of programmed cell death protein 1 (PD1) expression by HH7-2tg cells following co-culture with IFN-γ-treated colonic IEC organoids in the presence of the HH_1713 peptide (**Figures 4J and K**). This response was again dependent on pre-stimulation of the colonic IEC organoids with IFN-γ, as well as the presence of cognate antigen, and was inhibited by the addition of anti-MHC II blocking mAb (**Figure 4J**). Notably, we also detected significant upregulation of programmed death-ligand 1 (PD-L1 (*CD274*)) expression upon iCM stimulation of organoids (**Figure 1D**). These results highlight that although colonic IECs are able to activate MHC II-restricted HH7-2tg-FoxP3^hCD2^ cells in an antigen-specific manner, they may also render them susceptible to subsequent regulation via PD1.

**Figure 4:**
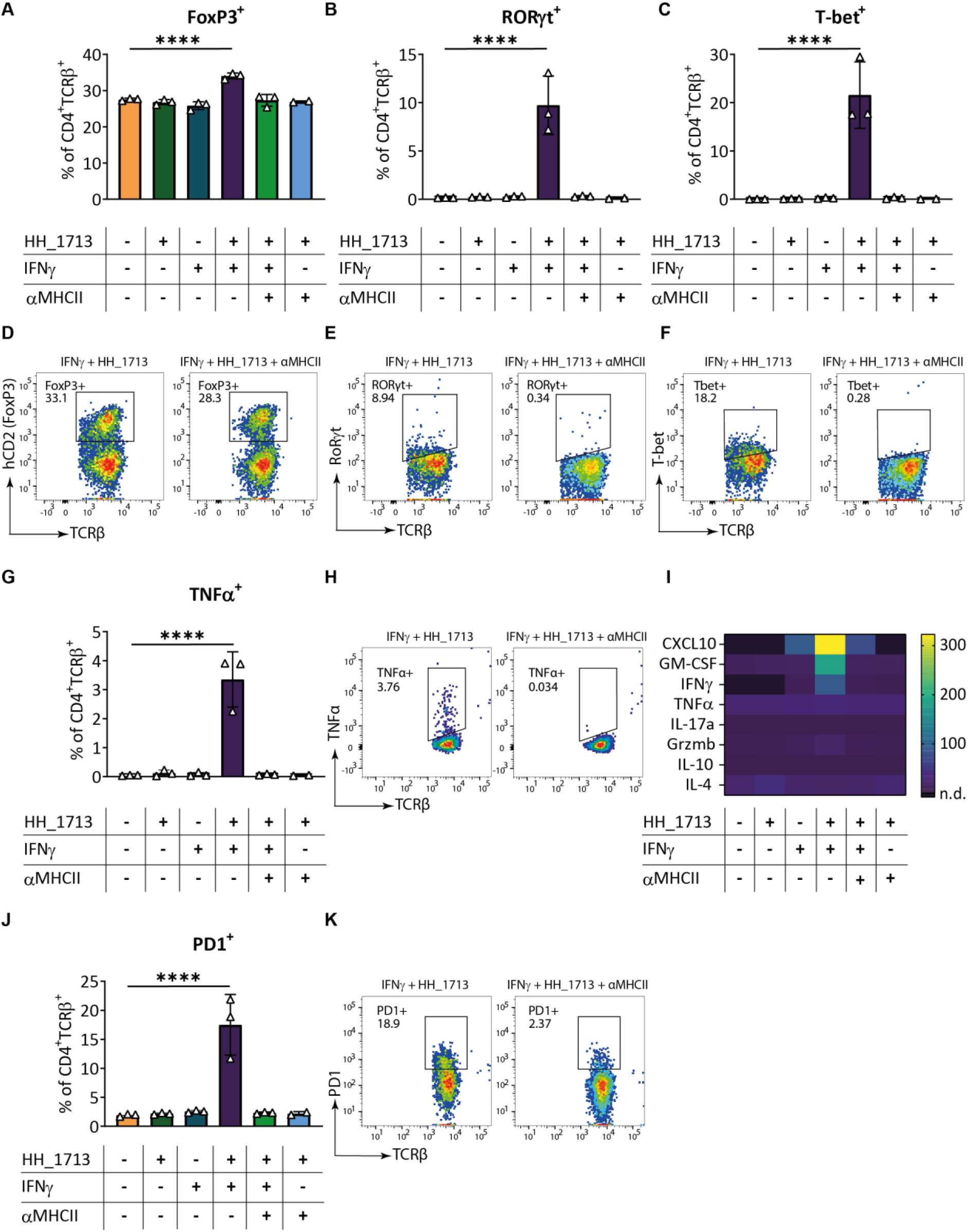
MHC II-expressing IECs induce antigen specific activation of CD4 T cells *ex vivo*. (A-K) 100000 pre-activated HH7-2tg-FoxP3^hCD2^ CD4 T cells were co-cultured with IEC^WT^ organoids for three days. (A-C) HH7-2tg-FoxP3^hCD2^ CD4 T cells were isolated and FoxP3 (A), RORγt (B) and T-bet (C) expression analysed by flow cytometry. (D-F) Representative FACS plots of FoxP3^+^ (D), RORγt^+^ (E) and T-bet^+^ (F) HH7-2tg-FoxP3^hCD2^ CD4 T cell frequencies, when co-cultured with colonic IEC^WT^ organoids pre-treated with IFN-γ (to upregulate MHC II expression) and peptide pulsed with HH_1713 on the left, and additional αMHCII treatment on the right. (G) HH7-2tg-FoxP3^hCD2^ CD4 T cells were isolated and TNFα expression analysed by flow cytometry. (H) Representative FACS plots of TNFα^+^ HH7-2tg-FoxP3^hCD2^ CD4 T cell frequencies, when co-cultured with colonic IEC^WT^ organoids pre-treated with IFN-γ (to upregulate MHC II expression) and peptide pulsed with HH_1713 on the left and additional αMHCII treatment on the right. (I) Supernatants of co-cultures were analysed by Luminex for CXCL10, GM-CSF, IFN-γ, TNFα, IL-17a, Grzmb, IL-10 and IL-4. (J) HH7-2tg-FoxP3^hCD2^ CD4 T cells were isolated and PD-1 expression analysed by flow cytometry. (K) Representative FACS plots of PD-1^+^ HH7-2tg-FoxP3^hCD2^ CD4 T cell frequencies, when co-cultured with colonic IEC^WT^ organoids pre-treated with IFNγ (to upregulate MHC II expression) and peptide pulsed with HH_1713 on the left and additional αMHCII treatment on the right. Representative data from one of three independent experiments, with three technical replicates for each co-culture condition, except for the HH_1713 + αMHCII condition, which consists of two technical replicates (A-H, J and K). Three technical replicates for each co-culture condition, except for the IFN-γ + HH_1713 and HH_1713 + αMHCII condition, which consists of four and two technical replicates, respectively (I). Symbols denote individual technical replicates. Bars represent mean (± SD). Statistical significance between the groups was determined by repeated measure mixed effects analysis with Tukey’s multiple comparisons test, with a single pooled variance. P-values of ctrl vs. HH_1713+IFN-γ conditions are shown. P-value summary: *p<0.05; **p<0.01; ***p<0.001; ****p<0.0001.

### Colonic IEC MHC II expression fine-tunes antigen-specific T cell responses in the inflamed intestine

To assess whether colonic IEC MHC II expression could shape antigen-specific CD4^+^ T cell responses in the context of intestinal inflammation *in vivo*, we treated IEC^ΔMHCII^ and IEC^WT^ littermates with *H. hepaticus* + anti-IL10R and adoptively transferred pre-activated HH7-2tg effector CD4^+^ T cells five days later. We analysed the transferred HH7-2tg T cells after a further seven days (**Figure 5A**). First, we confirmed that Hh+αIL10R treatment led to robust induction of colonic IEC MHC II expression in IEC^WT^ mice, but not in IEC^ΔMHCII^ littermates by flow cytometry (**Figure 5B and C**). At sacrifice, we detected comparable frequencies of HH7-2tg T cells in the lamina propria of IEC^ΔMHCII^ and IEC^WT^ recipients (**Figure 5D**), indicating similar levels of expansion and accumulation. However, when we examined their effector functions, we found significantly increased proportions of IFN-γ^+^ and a trend towards increased proportions of TNFα^+^ HH7-2tg T cells in IEC^ΔMHCII^ recipients compared to IEC^WT^ recipients (**Figure 5E and F**). Despite changes in antigen-specific effector functions, *H. hepaticus* colonisation levels were not significantly altered in IEC^ΔMHCII^ mice relative to their IEC^WT^ littermates (**Figure 5G**). Lipocalin-2, a faecal biomarker of intestinal inflammation^32^ revealed comparable levels of inflammation in IEC^ΔMHCII^ and IEC^WT^ mice (**Figure 5H**). Assessment of the histopathology of the caecum and colon revealed comparable disease severity between IEC^ΔMHCII^ and IEC^WT^ littermates (**Figure 5I-K**). Overall, these data indicate that the presence of functional MHC II on colonics IECs dampens the effector function of antigen specific CD4^+^ T cells in the inflamed intestine and that this fine-tuning acts to limit their production of pro-inflammatory cytokines.

**Figure 5:**
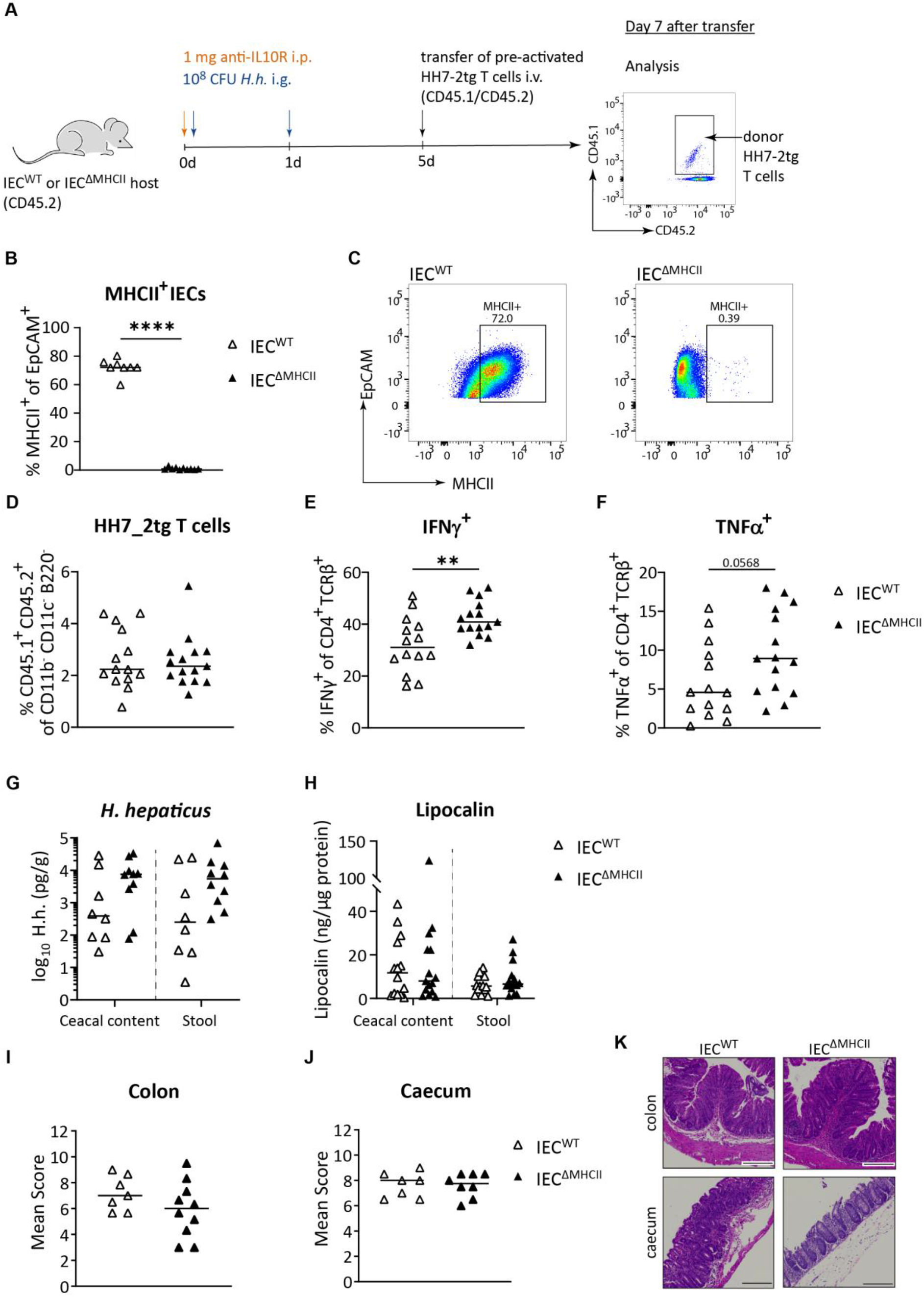
Colonic IEC MHC II expression fine-tunes antigen-specific T cell responses in the inflamed intestine. (A-K) FACS sorted naïve HH7-2tg CD4 T cells were pre-activated (3d anti-CD3/CD28 and RA), rested (2d RA and IL-2), and adoptively transferred into *H. hepaticus* colonised IEC^WT^ and IEC^ΔMHCII^ littermates. Mice were sacrificed 12d p.i. and tissues analysed for the presence of HH7-2tg CD4 T cells by flow cytometry. (A) Schematic of treatment protocol. (B) Colonic IEC MHC II expression of IEC^WT^ and IEC^ΔMHCII^ littermates was assessed by flow cytometry. (C) Representative FACS plots of MHCII^+^ IECs of IEC^WT^ and IEC^ΔMHCII^ littermates. (D) HH7-2tg CD4 T cell frequencies in the colonic lamina propria were assessed by flow cytometry. (E-F) LPLs were restimulated with PMA, Ionomycin and Golgi-Stop/Golgi-Plug for 3.5h. Frequencies of IFN-γ^+^ (E) and TNFα^+^ (F) HH7-2tg CD4 T cells were assessed by flow cytometry. (G) *H. hepaticus* DNA quantities were determined by qPCR in caecal content and stool of IEC^WT^ and IEC^ΔMHCII^ littermates. (H) Lipocalin quantities were determined by ELISA in caecal content and stool of IEC^WT^ and IEC^ΔMHCII^ littermates. (I and J) Histopathology of the colon (I) or caecum (J) was assessed on day 12. (K) Representative H&E sections of colon (top row) and caecum (bottom row) of IEC^WT^ and IEC^ΔMHCII^ littermates at day 12. Data from two pooled independent experiments with total mouse numbers per group as follows; n=15 IEC^ΔMHCII^ and n=14 IEC^WT^ treated mice (A-F, H). Data are representative of one of two independent experiments with total mouse numbers per group as follows; n=10 IEC^ΔMHCII^ and n=8 IEC^WT^ treated mice (G, I and J). Symbols denote individual mice. Horizontal bars indicate group medians. Statistical significance was determined using Mann-Whitney test (B-F and I-J) or two-way ANOVA with Sidak’s multiple comparisons test (G and H). P-value summary: *p<0.05; **p<0.01; ***p<0.001; ****p<0.0001. Scale bars, 200*μ*m. *H.h*. – *Helicobacter hepaticus*.

## Discussion

Since the 1970s, when MHC II expression by IECs during intestinal inflammation was first described^6–8^, researchers have debated the potential role of antigen presentation by IECs in IBD pathogenesis^13,17,18^. Although modern cellular and genetic techniques have allowed greater insight, there is still no consensus as to the primary functional role of MHC II expression by IECs, especially in the context of intestinal infection and inflammation. To investigate this, we evaluated IEC-specific ablation of MHC II in well characterized mouse models of bacterially-triggered intestinal inflammation in combination with *ex vivo* IEC organoid cultures and antigen-specific CD4^+^ T cells. Our data indicates that presentation of MHC II-peptide complexes by IECs fine-tunes the CD4^+^ T cell response, limiting their inflammatory potential. This is important because it suggests that local MHC II antigen presentation by IECs may play a role in limiting inflammatory damage in the epithelial layer, allowing restoration and maintenance of barrier function.

Our study of colonic organoids exposed to the inflammatory milieu demonstrated antigen processing and presentation as a key epithelial response to inflammation. The IFN-γ signalling pathway was also highly upregulated, and, consistent with previous reports^16^, we observed that IFN-γ is a dominant inflammatory mediator that induces MHC II expression in IECs, through activation of CIITA. The identities of the antigens presented by IECs in the inflamed colon are not known, but our organoid analyses indicated that IECs can take up and process exogenous antigen and that they can present cognate peptide to antigen-specific CD4^+^ effector T cells.

Our IEC organoid – T cell co-culture experiments showed that MHC II antigen presentation by colonic IEC organoids enhanced the proportions of effector T cells expressing molecules associated with pathogenic CD4^+^ T cells, such as T-bet, RORγt, TNFα, IFN-γ and GM-CSF. These results suggest that MHC II^+^ colonic IECs can provide activation and/or survival signals to effector CD4^+^ T cells, and these interactions may result in cytokine secretion toward the epithelium. These observations echo previous reports of antigen specific activation and induction of CD4^+^ T cell effector functions by IEC MHC II expression *ex vivo*^18,33^. Convincing evidence that IEC MHC II expression can drive pathogenic CD4^+^ Th1 cell responses *in vivo* was provided by a recent study of experimental GvHD, where small intestinal IEC MHC II expression was shown to induce acute GvHD small intestinal pathology and lethality^16^. Concomitantly with increased cytokine production by T cells interacting with IEC organoids, we also observed upregulation of PD-1, suggesting that these cells may be more susceptible to regulation through the PD-1/PDL-1 pathway *in vivo*. In addition, we saw small but consistent increases in the proportions of Tregs in our *in vitro* cultures. This led us to test whether the enhanced effector T cell activation seen in our *in vitro* analysis and during the unusual context of GvHD would be recapitulated during bacteria-induced inflammatory responses *in vivo*.

The IEC^ΔMHCII^ mice allowed us to interrogate the potential impact of MHC II expression by IECs on the outcome of bacterially-triggered intestinal inflammation *in vivo*. We used the A/E pathogen *C. rodentium* as a model of acute colitis and *H. hepaticus* infection with anti-IL10R treatment as a model of chronic colitis. Both models showed comparable levels of inflammation in IEC^ΔMHCII^ and IEC^WT^ littermates, indicating that MHCII expression by IECs is dispensable for bacterially-driven colitis. During *C. rodentium* infection, IEC^ΔMHCII^ mice had comparable levels of intestinal inflammation, transient weight loss and pathogen colonization to their WT littermates and did not exhibit any morbidity or mortality. Our data contrast starkly with a recent study which reported that IEC^ΔMHCII^ mice exhibited more severe intestinal pathology, increased pathogen loads and 50% mortality following *C. rodentium* infection, which correlated with impaired production of secretory IgA^34^. Although many factors could contribute to these discrepancies, high mortality following infection with a pathogen that normally elicits a mild, self-limiting disease suggests fundamental biological differences. Some Villin-Cre transgenic lines can exhibit a low level of Cre recombinase activity in extraintestinal tissues which can lead to germline deletion of the floxed alleles^28^. Indeed, we observed this in multiple strains we generated using Villin-Cre mice, including the MHC II I-Ab ‘floxed’ strain used in this study. Variable proportions of the colony were determined to be complete knockouts for MHC II (data not shown). This possibility means that validation of IEC-specific deletion should be performed for every individual mouse, as we did in this study. It is not clear whether the conflicting study^34^ performed individual validation to circumvent this issue, but it is notable that mice lacking secretory IgA do not succumb to *C. rodentium* infection^35,36^, whereas complete MHC II KO mice do^37^. Furthermore, although IFN-γ^-/-^ mice are unable to upregulate MHC II on colonic IECs during *C. rodentium* infection, they did not exhibit any mortality^38^. Taken together, these results provide a plausible explanation for differing results and support our conclusions that colonic IEC MHC II expression has little impact on *C. rodentium* induced intestinal inflammation and is not required for survival from this A/E pathogen.

Although we found that in wild type mice almost all colonic IECs expressed MHC II following Hh+anti-10R treatment, we observed no significant differences in intestinal pathology between IEC^ΔMHCII^ mice and their wild type littermates. These results demonstrate that IEC MHC II expression is dispensable for the induction of intestinal pathology in this model. Together, these results emphasise that IEC MHC II expression is not required for the induction of colitogenic CD4^+^ T effector responses in the intestine. We also found no major change in the proportions of effector or regulatory CD4^+^ T cell subsets in the lamina propria of IEC^ΔMHCII^ mice and their wild type littermates following Hh+anti-10R treatment, again contrasting with our *in vitro* data.

Despite these conclusive *in vivo* results, the semi-quantitative nature of the histopathological scores used to assess Hh+anti-10R-driven colitis, together with minor inter-experimental variations typical of *in vivo* models, makes them rather insensitive as a means of monitoring more subtle changes in disease. In addition, the severe inflammatory response comprises a polyclonal CD4^+^ T cell response directed against antigens from both *H. hepaticus* and other microbiota components. This prompted us to use an adoptive transfer system to assess the effects of colonic IEC MHC II expression on *H. hepaticus-specific* CD4^+^ effector T cells during Hh+anti-10R-driven colitis. These experiments revealed that, even in the midst of severe intestinal inflammation, IEC MHC II expression acts to attenuate the inflammatory potential of *H. hepaticus-specific* CD4^+^ effector T cells.

The inhibitory effects were observed on *H. hepaticus-specific* CD4^+^ effector T cells isolated from the intestinal lamina propria, which raises the question of how these T cells can be influenced by MHC II molecules expressed on IECs. There are several possibilities, including direct inhibition of effector CD4^+^ T cells through interactions with IECs, IEC-T cell interactions at the expense of interactions with more inflammatory APCs, or tuning of other immune cells that interact with CD4 T cells^39^. It was shown that epithelial histone deacetylase 3 is essential for NFκB-dependent regulation of epithelial MHC II, which dampens the local accumulation of commensal-specific Th17 cells^40^. Our organoid co-culture experiments indicate that MHC II expressing IECs can directly modulate CD4^+^ effector T cells, and we also observed secretion of the Th1-recruiting, pro-inflammatory chemokine CXCL10 in these co-cultures, consistent with previous studies in murine and human IEC organoids^41,42^. This highlights a potential positive feedback loop through which IECs may recruit further effector CD4^+^ T cells into the epithelium. It is notable that CD4^+^ T cell effector cytokines have been reported to regulate the renewal (IL-10) and differentiation (IFN-γ, IL-17A and IL-13) of Lgr5+ intestinal stem cells^15^, therefore our results are compatible with the concept that IEC-CD4^+^ T cell crosstalk regulates epithelial barrier dynamics and integrity during intestinal inflammation. As our co-culture data suggests, interaction with IECs may render effector CD4^+^ T cells susceptible to regulation through the PD-1/PD-L1 pathway. Our data supports the hypothesis that IECs directly inhibit effector CD4^+^ T cells through PD-1/PD-L1 because we found that PD-L1 was one of the most highly upregulated genes in colonic IEC organoids that were exposed to inflammatory conditioned medium (iCM) and IECs from IBD patients were described to express PD-L1 protein^43^. Further work to understand the effect of local T cell accumulation on the epithelium itself may further clarify the “purpose” of MHCII expression on epithelial cells in the context of inflammation.

The data presented in this study along with the growing body of work on antigen presentation by non-conventional APCs demonstrate the complexity of immune responses at barrier tissues. Here we demonstrate inhibitory tuning of the CD4 T cell response by epithelial MHCII, but it is not entirely surprising that IEC MHC II expression may also boost effector CD4^+^ T cell responses under certain circumstances, as there are several enteric pathogens, such as rotavirus^44^, adenovirus^45^, *Salmonella typhimurium*^46^ and *Cryptosporidium tyzzeri*^47^, that can directly infect small intestinal IECs. Future studies should assess the role of epithelial MHC II expression on pathogen clearance and pathology during infection with these enteric pathogens. MHC II expression by the epithelium is one important tool the intestine uses to maintain homeostasis, and further work will be needed to define additional direct and indirect mechanisms of immune control by IECs.

## Methods

### Mice

H2-Ab1^fl/fl^ mice were bought from JAX laboratory (B6.129X1-H2-Ab1tm1Koni/J).^26^ H2-Ab1^fl/fl^ mice were crossed to B6.Cg-tg(VIL-1-cre)997Gum/J mice^27^ (JAX laboratory) to generate H2-Ab1^VC^ and H2-Ab1^fl/fl^ mice. HH7-2tg TCR transgenic mice were kindly provided by Dr. Dan R. Littman (New York). HH7-2tg TCR transgenic mice were crossed to B6.Foxp3^hCD2^ IL-10^GFP^ reporter mice (HH7-2tg-FoxP3^hCD2^)^48^. The strains were maintained in accredited animal facilities at the University of Oxford, UK. Mice were bred under specific pathogen-free conditions with environmental enrichment and routinely screened for pathogens and *Helicobacter* species. All experiments were performed in accordance with the UK Scientific Procedures Act of 1986 and under Project Licenses (PP9127884 and 30/3423) authorised by the UK Home Office Animal Procedures Committee and approved by the Sir William Dunn School of Pathology, and Kennedy Institute of Rheumatology ethical review committee.

Mice were bred hemizygous for the Cre allele, resulting in H2-Ab1^fl/fl^ Cre-negative and H2-Ab1^fl/fl^ Cre-positive (H2-Ab1^VC^) littermates. Both female and male, age- and sex-matched littermates were used and co-housed throughout the experiment. Mice were analysed at 6-20 weeks of age.

### Colitis models

*H. hepaticus* (NCI-Frederick isolate 1A^49^) was cultured in TSB media supplemented with 10% FCS and trimethoprim (5μg/ml), vancomycin (10μg/ml) and polymyxin B (25IU/ml) (TVP, Oxoid) under microaerophilic conditions (10% CO_2_, 10% H_2_, 80% N_2_)^50^. Chronic colitis was induced by infecting mice intragastrically with 10^8^ CFU *H. hepaticus* on two or three consecutive days and injecting intraperitoneally 1mg of mAb anti-IL10R (clone 1B1.2) weekly. *C. rodentium* (ICC169) was grown in Luria Broth supplemented with nalidixic acid in log phase to an OD of 1. Acute colitis was induced by infecting mice intragastrically with 10^9^ CFU *C. rodentium*^50^.

### Histological Assessment of Intestinal Inflammation

Mice were sacrificed at indicated time points. Tissue sections were fixed in buffered 10% formalin over-night and paraffin-embedded. Histological analysis of H&E stained proximal, middle and distal colon sections as well as caecum sections for intestinal inflammation was performed as previously described^50^. Briefly, inflammation was graded semiquantitatively on a scale from 0 to 3, for four or five criteria; (a) epithelial hyperplasia and goblet cell depletion, (b) lamina propria leukocyte infiltration, (c) area of tissue affected, (d) markers of severe inflammation, including crypt abscesses, sub-mucosal inflammation, and ulceration and (e) submucosal markers, including oedema, cell infiltration and follicle enlargement. Scores for criteria (a) to (d) were added up for an overall inflammation score between 0 and 12 for *H. hepaticus* + anti-IL10R colitis. Scores for criteria (a) to (e) were added up for an overall inflammation score between 0 and 15 for *C. rodentium* colitis. Data from the three colon regions were averaged to give an overall colon score. Scoring was performed by two scientists independently in a blinded fashion.

### HH7-2tg T cell transfer model

Splenic naïve HH7-2tg T cells were FACS sorted as B220^-^CD11c^-^CD11b^-^CD45^+^CD25^-^CD44^lo^CD62L^hi^CD4^+^TCRVβ6^+^. Naïve HH7-2tg cells were preactivated with anti-CD3 (5μg/ml) and anti-CD28 (2μg/ml) and retinoic acid (10nM) for 3d and rested with recombinant IL-2 (5ng/ml) and retinoic acid (10nM) for a further 3d. Dead cells were removed with the Miltenyi dead cell removal kit according to the manufacturer’s instructions. 700000 or 250000 HH7-2tg T cells were intravenously injected 5d after the induction of chronic colitis with *H. hepaticus* + anti-IL10R treatment. HH7-2tg T cells were analysed on day 7 after transfer.

### Isolation of IECs and lamina propria leukocytes

Colon and caecum were harvested from mice, contents removed and opened longitudinally. Tissue was washed, cut into 5mm pieces and placed in ice-cold PBS/0.1%BSA. The tissue was incubated 2 times in RPMI 1640/5%FCS supplemented with 5mM EDTA (Gibco) for 20min each at 37°C and 200rpm. The supernatant, containing IECs, was filtered through a 100μm cell-strainer and used for subsequent flow-cytometry analysis. The tissue was incubated in RPMI/5%FCS/10mMHEPES for 5min before being digested in 10ml RPMI/5% FCS/10mM HEPES supplemented with 0.4mg/ml type VIII collagenase (Sigma Aldrich, UK) and 40μg/ml DNase I (Roche, UK) for 45 - 60min at 37°C, 180rpm. Supernatants were filtered through a 70μm nylon mesh and the collagenase activity neutralised by adding EDTA at 5mM. After centrifugation cells were resuspended in 37.5% Percoll-solution (GE Healthcare) and centrifuged for 5min at 1800rpm. The cell pellet was resuspended in PBS/0.1%BSA/EDTA and used for subsequent flow-cytometry analysis.

### Antibodies, intracellular staining and flow cytometry

The following monoclonal antibodies were purchased from eBiosciences, BD Biosciences or BioLegend: CD11c (N418), CD11b (M1/70), B220 (RA3-6B2), CD4 (RM4-5), TCRβ (H57-597), TCR Vβ6 (RR4-7), CD25 (Ebio3c7), CD62L (MEL-14), CD44 (IM7), CD45.1 (A20), CD45.2 (104), CD45 (30-F11), FOXP3 (FJK-16s), GATA3 (TWAJ), RORγt (B2D or Q31-378), T-bet (eBio4B10), IL-17A (eBio17B7), TNFα (MP6-XT22), IFNγ (XM61.2), EpCAM (G8.8) and MHCII (M5/114.15.2). Dead cells were labelled using efluor-780 fixable-viability dye (eBioscience).

For assessment of transcription factors, cells were stained for surface markers, fixed and permeabilised, and stained for nuclear factors according to the manufacturer’s protocol (FOXP3 staining buffer set (eBioscience)). For cytokine analysis, cells were incubated for 3.5h in complete RPMI (10% FBS, 1x GlutaMax, 1x Pen/Strep, 10mM HEPES, 1x Sodium-pyruvate, 55uM beta-mercaptoethanol) and stimulated with phorbol 12-myristate 13-acetate (PMA) (50ng/ml; Sigma), ionomycin (1ug/ml; Sigma) and GolgiStop/Golgi-Plug (1:1000, BD Biosciences). Cells were stained for surface markers, fixed and permeabilised, and then subjected to intracellular cytokine staining according to the manufacturer’s protocol (Cytofix/Cytoperm buffer set (BD Biosciences)).

Samples were acquired on LSR II or Fortessa X20 flow cytometers, cell sorting was performed using the Aria II (all BD Biosciences) and analysed using FlowJo software (Tree Star).

### Organoid Culture

Organoids were cultured as previously described^51^. Colonic crypts were isolated from longitudinal opened and cleaned colon tissue of H2-Ab1^fl/fl^ mice or H2-Ab1^VC^ mice by 2mM EDTA in PBS incubation for 90min at 4°C. The tissue was vigorously resuspended in PBS/10% FCS and the supernatant containing the crypts filtered through a 100μm cell strainer and pelleted. Crypts were seeded in 20μl Matrigel/well (Corning) in a 24-well plate. After polymerisation of the Matrigel, 400ul of complete organoid medium was added [Advanced Dulbecco’s Modified Eagle Medium (DMEM)/F12 supplemented with 1% Pen/Strep, 10mM HEPES, 1X GlutaMax, 50ng/ml mEGF, 1× N2, 1× B27 (all from Invitrogen), 1mM N-acetylcysteine (NAC), 10μM RhoK-inhibitor (Y-27632 dihydrochloride monohydrate) (both from Sigma) and conditioned medium containing Wnt3a (20%, or 330pM recombinant Wnt3a (from ImmunoPrecise)), Noggin (20%, or 1% recombinant Noggin (from ImmunoPrecise)) or R-spondin (10%)]. The conditioned media were generated from Rspol-Fc Hek 293T cells (kind gift from Dr. Calvin Kuo, Stanford) and Nog-Fc Hek 293T cells (kind gift from Dr. Gjis van den Brink, Amsterdam) and Wnt3A L cells (ATCC CRL-2647). The medium was changed every 2-3d and organoids were expanded over several passages before they were used for experiments with indicated stimuli.

### BMDC culture

Bone marrow cells of the tibia and femur were cultured in 10ml of complete RPMI medium (10% FBS, 1x Pen/Strep, 1x Sodium-Pyruvate, 25mM HEPES and 55uM beta-mercaptoethanol) and GM-CSF (20ng/ml, Peprotech). 10ml of new medium was added with GM-CSF (20ng/ml) on day 3 and 6. On day 9-12, non- and loosely-adherent cells were harvested and used for experiments.

### DQ Ova assay

BMDCs were seeded at 1*10^5 cells/well in a flat-bottom 96-well plate prior to the experiment. Organoids were treated with IFN-γ (20ng/ml) for 16h prior to the experiment (where indicated). BMDCs and organoids were cultured with 10μg/ml DQ Ovalbumin (Thermo Fisher Scientific) at 4°C, 37°C or with no DQ Ovalbumin for the indicated time, washed and analysed using flow cytometry.

### Organoid-CD4^+^ T cell co-culture

Splenic naïve HH7-2tg-FoxP3^hCD2^ T cells were FACS sorted, pre-activated for 3d (anti-CD3 (5μg/ml) + anti-CD28 (2μg/ml)) and rested for 2d with recombinant IL-2 supplementation (5ng/ml). Colonic organoids were split 1:1 2 days prior to IFN-γ stimulation and filtered using a 100μm cell-strainer. Depending on the experimental condition, organoids were stimulated with IFN-γ (20ng/ml), IFN-γ + anti-MHCII (1μg/ml) or left untreated for 16h prior to the start of the co-culture. At the start of the co-culture, organoids were isolated from the Matrigel using Cell-recovery solution (Corning) for 20min at 4°C. Pre-activated HH7-2tg-FoxP3^hCD2^ T cells were subjected to dead cell removal using the Miltenyi Dead cell removal kit, according to the manufacturer’s recommendations. 100000 HH7-2tg-FoxP3^hCD2^ T cells were co-cultured with colonic organoids from 2 wells of a 24-well plate in 30μl of collagen/Matrigel mix (PureColl/Matrigel, 10xMEM, 7.5% sodium bicarbonate, 1 ug/ml CCL19). HH7-2tg T cells and colonic organoids were co-culture for 2d, depending on the condition with 1ug/ml HH_1713 and/or 1ug/ml anti-MHCII. Supernatant was removed and analysed by Luminex analysis. HH7-2tg-FoxP3^hCD2^ T cells were isolated using collagenase VII (100U/ml, Sigma Aldrich) for 1h at 37°C and subjected to flow cytometric analysis. For cytokine expression analysis, HH7-2tg-FoxP3^hCD2^ T cells were incubated with Golgi-Plug and Golgi-Stop (1:100, BD Biosciences) for 4h at 37°C.

### Luminex analysis

Supernatants of organoid HH7-2tg-FoxP3^hCD2^ T cells cultures were subjected to Luminex analysis. The following targets (CXCL10, Granzyme-B, IL-2, IL-10, TNFα, GMCSF, IFN-γ, IL-4 and IL-17A) were measured according to the manufacturer’s instructions (Biotechne) and analysed with the Bio-Rad Bio-Plex 20 system.

### RNA and qPCR analysis

Tissues were disrupted using lysis beads (Precellys) and a TissueLyserII (Qiagen) in RLT buffer (Qiagen). Cells were lysed in RLT (with 143mM beta-mercaptoethanol) and RNA purified using the RNeasy kit (Qiagen). 0.8-2μg of RNA were reverse transcribed using the High-Capacity cDNA reverse transcription kit and quantitative real-time PCR was carried out with the ViiA 7 Real-time PCR system (both Applied Biosystems). qPCR Mastermix (PrimerDesign) was used for Taqman reactions. Taqman probes for *H2-Ab1* (Mm00439216_m1), *Ciita* (Mm00482914_m1) and *Hprt* (Mm03024075_m1) were from Applied Biosystems and primers for *H. hepaticus* (forward: CCG CAA ATT GCA GCA ATA CTT; reverse: TCG TCC AAA ATG CAC AGG TG) from Merck. Relative expression was analysed according to the Pfaffl method^52^.

### Immunofluorescent imaging

Fresh frozen tissue sections were permeabilised and blocked using 0.5% Saponin and 10% serum from the same species as secondary antibodies. Anti-MHCII (clone M5/114.15.2, invitrogen), anti-CD3 (Abcam), anti-Ecadherin (clone 36/E-cadherin, BD Biosciences) (all 1:100; in 5%BSA/PBS) were incubated for 1h at RT. Tissue was washed and incubated with secondary antibodies AlexaFluor488, AlexaFluor555 and AlexaFluor647 (all 1:400; in 5% BSA/PBS (lifetechnologies)) for 30min at RT. Tissue was washed, incubated with DAPI for 5min at RT (1:5000, in PBS (Sigma Aldrich)) and washed. Sections were mounted using ProLong Gold Antifade (Invitrogen).

Organoids were grown on coverslips. Following 16h of IFN-γ stimulation cells were fixed with 2% paraformaldehyde at room temperature followed by methanol treatment at −20°C. Cells were blocked in PBS/5% BSA/5% goat serum/0.5% saponin, followed by staining with pAb mouse-anti-E-cadherin (BD Biosciences), rat-anti MHCII (M5/114.15.2, Biolegend), rabbit-anti-CD3 (R&D) and goat-anti-mouse Alexa Fluor 488, goat-anti-rat Alexa Fluor 555, goat-anti-rabbit Alexa Fluor 647 (lifetechnologies). Images were acquired with an Olympus Fluoview FV1000 confocal microscope or Zeiss980 confocal microscope and ImageJ Software.

### Generation of Inflammatory Conditioned Medium (iCM)

LPLs were isolated from C57BL/6 mice at 2 weeks of *H. hepaticus* + anti-IL10R colitis and 2.3×10^6^ cells/ml were seeded in complete RPMI for 20h. Supernatants from cells of 7 mice were pooled and frozen at −80°C.

### WesternBlot

For immunoblot analysis cells were lysed after indicated times in RIPA buffer and 3.7μg total protein were analysed per lane. For detection the following primary antibodies were used: phospho-Stat3, phospho-Stat1, phospho-ERK, phospho-p65, phospho-p38, phospho-JNK, phospho-AKT and GAPDH (all Cell Signalling).

### RNAseq analysis

Colonic organoids were seeded for stimulation at p4, p6 and p7. 24h prior to stimulation full medium was reduced to organoid medium containing 1% Wnt3a. Colonic organoids were stimulated with 10% cCM or iCM for 24h prior to lysis and RNA isolation (Qiagen RNeasy). Quadruplicates were sequenced.

Poly-A+, stranded RNA-seq libraries were prepared based on the Illumina (Illumina, CA) TruSeq Stranded mRNA sample prep kit using random hexamer priming. Resulting libraries were sequenced using the Illumina Hiseq2500 rapid run (100bp PE) with the Agilent 2100 Bioanalyzer Desktop System.

Raw reads were assessed for quality using Fastqc (v0.11.7, https://www.bioinformatics.babraham.ac.uk/projects/fastqc/). Paired reads were aligned to the mouse genome (build mm10) using HISAT2 (v2.1.0) [PMID: **31375807**]. Gene counts were produced using featureCounts [PMID: **24227677**] (v1.6.0) against Ensembl genesets (Ensembl build 91 [PMID: **30407521**]). Genes were removed from downstream analysis if they contained fewer than 10 reads in more than 4 samples. Differential expression analysis on cCM vs iCM treated organoids was performed using DESeq2 [PMID: **25516281**] and genes were considered significant at an adjusted p-value < 0.05.

### Gene set enrichment analysis

We ranked genes according to both significance and fold change (i.e. −log10(padj) x log2(fold change)) for input into Fast Gene Set Enrichment Anlaysis (fgsea [https://www.biorxiv.org/content/10.1101/060012v1]). We used mouse canonical pathways from MsigDb (https://www.gsea-msigdb.org/gsea/msigdb/mouse_geneset_resources.jsp, m2.cp.v0.3) as the input gene sets. Fgsea was run against gene sets that had a minimum of 15 genes and a maximum of 500 genes. We considered gene sets significantly enriched at an adjusted p-value < 0.05.

## Quantification and statistical analysis

The statistical test applied and the number of experimental repeats are specified in the figure legends. Differences were considered statistically significant when p < 0.05 (*p < 0.05, **p < 0.01, ***p < 0.001). Statistics were calculated using GraphPad Prism 9 software.

## Data Availability

The accession number for the raw RNA sequencing data reported in this paper is E-MTAB-12414.

## Acknowledgments

We thank members of Kevin Maloy’s and Fiona Powrie’s groups for helpful discussions and technical help. We are grateful to Gijs van den Brink for providing Noggin- and R-spondin-producing cell. We further thank the following Kennedy Institute and Sir William Dunn School of Pathology core facility members for excellent technical support and training: J. Webber, M. Maj, B. Stott, I. Parisi, R. Cook, C. Lagerholm and V. Nechyporuk-Zloy. K.J.M and J.P. were funded by the MRC Project Grant MR/N02379X/1. K.J.M. is supported by a Wellcome Trust Investigator award (102972). C.H. was supported by a Graduate Student Scholarship from the MRC (MR/R502224/1).

## Author Contributions

C.E.H., E.T., J.P., F.M.P. and K.J.M. conceptualised the study. C.E.H., A.J, A.B., Y.G., M.P., M.F., E.M., C.P. performed experiments. C.E.H., N.I., S.P. and J.P. analysed data. J.P. and K.J.M. secured funding. C.E.H., E.T. and K.M. wrote the manuscript. All authors agreed to submit the manuscript, have read and approved the final draft, and take full responsibility of its content, including the accuracy of the data and their statistical analysis.

## Abbreviations

(IECs): Intestinal epithelial cells
(MHC II): Major histocompatibility complex class II
(IBD): Inflammatory Bowel Disease
(GvHD): Graft-versus-host disease
(Tregs): Regulatory T cells
(A/E): Attaching-effacing
(LPLs): Lamina propria leucocytes
(iCM): Inflammatory conditioned medium
(cCM): Control conditioned medium
(RNA-Seq): RNA sequencing
(APC): Antigen presenting cell
(CIITA): class II major histocompatibility complex transactivator
(*H. hepaticus*): *Helicobacter hepaticus*
(*C. rodentium*): *Citrobacter rodentium*
(HH7-2tg cells): *H. hepaticus-specific* CD4^+^ T cells
(TF): Transcription factors
(PD1): Programmed cell death protein 1
(PD-L1): Programmed death-ligand 1
(MFI): Median fluorescent intensity
(CFU): Colony-forming units
(H&E): Hematoxylin and eosin

