## Supplementary Figures for "MHC class II antigen presentation by intestinal epithelial cells fine-tunes bacteria-reactive CD4 T cell responses"

### Supplemental figure S1

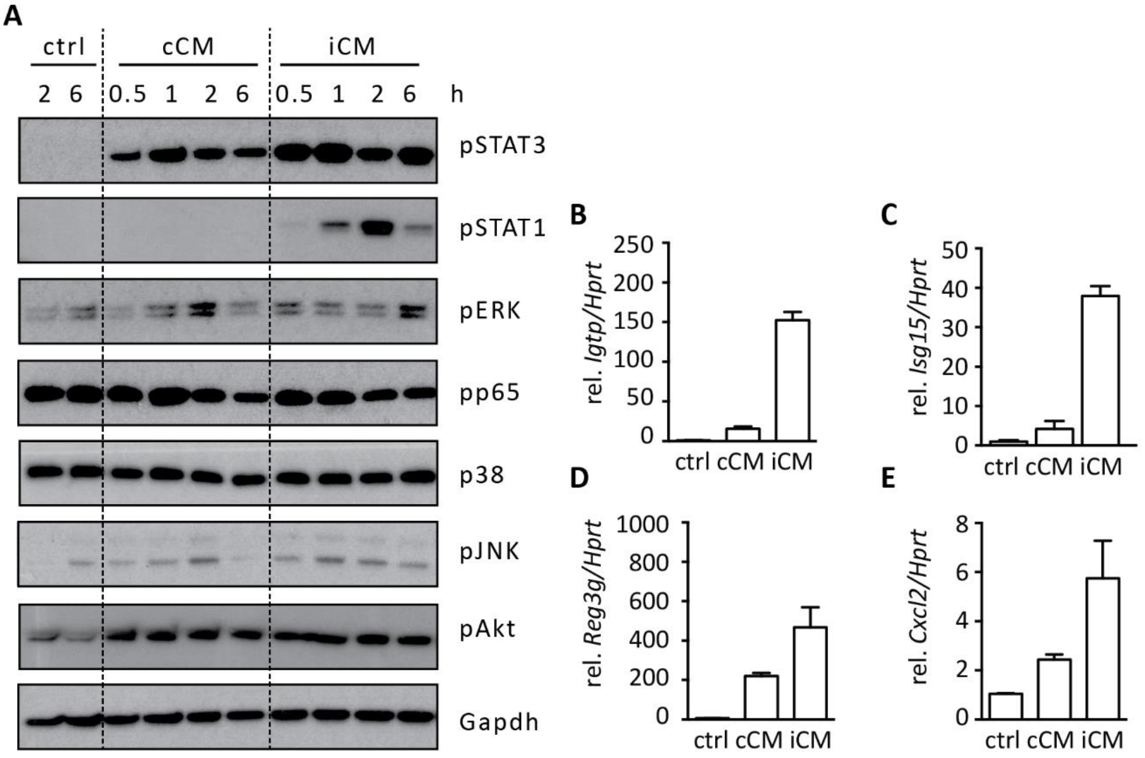


**Supplementary Figure 1: Related to Figure 1. Colonic IEC organoids respond to iCM stimulation with the rapid phosphorylation of STAT1 and 3**

(A-E) Colonic IECs were grown *ex vivo* in organoids. iCM was harvested from LPL cultures from colitic mice (H.h.+αIL10R) and cCM was harvested from LPL cultures from untreated mice.

(A) Colonic organoids were stimulated with 10% iCM or cCM for indicated times and lysates were probed with antibodies directed against indicated proteins.

(B-E) qPCR analysis of *Igtb* (B), *Isg15* (C), *Reg3g* (D) and *Cxcl2* (E) expression by colonic organoids treated with 10% iCM for 20h.

Data are representative of one experiment. Data are shown as means (± SEM). Statistical significance was determined using Mann-Whitney test. iCM – inflammatory conditioned medium; cCM – control conditioned medium, ctrl – control.

**Supplemental figure S2**

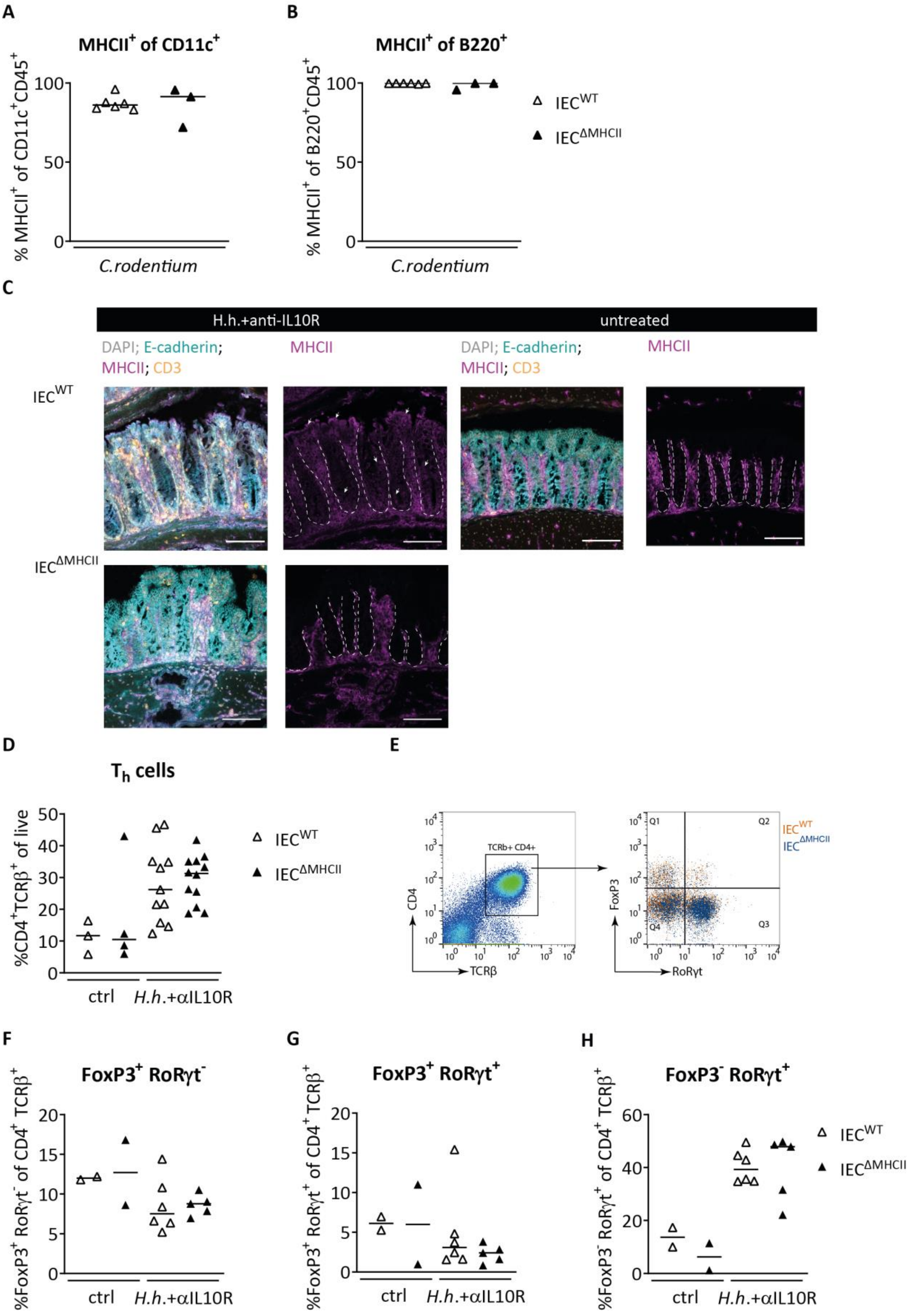

**Supplementary Figure 2: Related to Figure 2 and 3. Lack of IEC MHC II expression does not impact FoxP3 and RORγt expression by CD4 T cells during chronic colitis**

(A and B) $IEC^{WT}$ and $IEC^{\Delta MHCII}$ littermates were orally infected with $10^9$ CFU *C. rodentium*.

(A and B) Frequencies of $MHCII^+$ $CD11c^+$ (A) and $B220^+$ (B) cells were analysed by flow cytometry at d14.

(C-H) $IEC^{WT}$ and $IEC^{\Delta MHCII}$ littermates were orally infected with $10^8$ CFU *H. hepaticus* on three consecutive days and i.p. injected with 1mg anti-IL-10R weekly.

(C) MHC II (magenta), CD3 (orange) and E-cadherin (turquoise) stainings of colon sections isolated at day 14 after colitis induction of *H.h.*+αIL10R treated or untreated mice. Counterstaining with DAPI (grey).

(D) Frequencies of lamina propria $CD4^+$ $TCR\beta^+$ T cells were analysed by flow cytometry on day 28.

(E) Schematic of FACS gating strategy.

(F-H) Frequencies of $FoxP3^+$ $RoR\gamma t^-$ (F), $FoxP3^+$ $RoR\gamma t^+$ (G) and $FoxP3^-$ $RoR\gamma t^+$ (H) CD4 T cell subsets were analysed by flow cytometry on day 28.

Data from one experiment of n=3 $IEC^{\Delta MHCII}$ and n=6 $IEC^{WT}$ infected mice (A and B). Data from two pooled independent experiments with total mouse numbers per group as follows; n=12 $IEC^{\Delta MHCII}$ and n=11 $IEC^{WT}$ treated mice and n=4 $IEC^{\Delta MHCII}$ and n=3 $IEC^{WT}$ untreated mice (D). Data from one experiment with total mouse numbers per group as follows; n=5 $IEC^{\Delta MHCII}$ and n=6 $IEC^{WT}$ treated mice and n=2 $IEC^{\Delta MHCII}$ and n=2 $IEC^{WT}$ untreated mice (F-H). Symbols denote individual mice. Horizontal bars indicate group medians. Statistical significance between the groups was determined by Mann-Whitney (A and B) or two-way ANOVA with Tukey's multiple comparison test (D-H). Dashed line indicates intestinal epithelial crypt region. Arrows show examples of $MHCII^+$ IECs. Scale bars, $\mu$m.
